## supplemental figures and tables for "Rice stripe virus utilizes a *Laodelphax striatellus* salivary carbonic anhydrase to facilitate plant infection by direct molecular interaction"

#### Figure S1

A

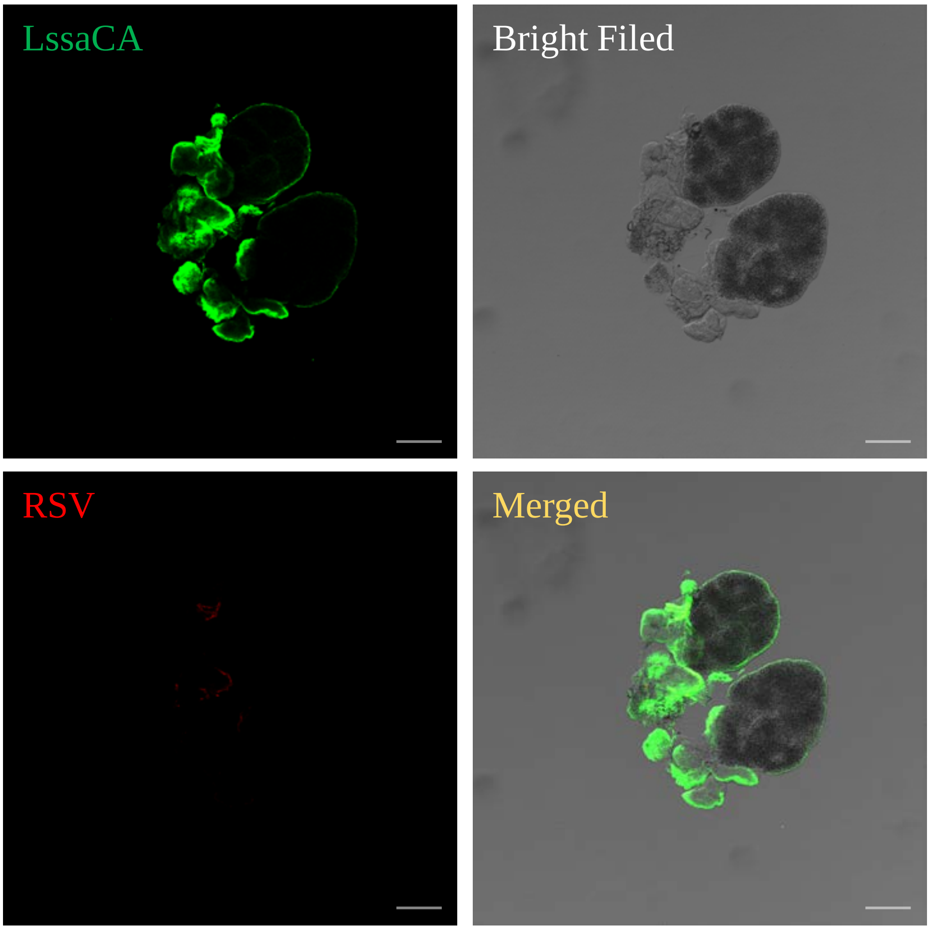


**Figure S1. Immunofluorescence localization of RSV and LssaCA in uninfected *L. striatellus* salivary glands.**

RSV (red) is absent in uninfected salivary glands, while LssaCA (green) is detected. Scale bar: 20 μm. Abbreviations: psg, principal salivary gland; asg, accessory salivary gland.

#### Figure S2

**A**

```
  1 MELLFHISMI LTIMAVSLAG MGRSSSFLRN LGGGRLGGPM APGEPEFPEV HEGGSSGFPW
 61 TPRSQYKSPQ QSPIDICTCN CKAMNYRPLR VNYAKTDRRD GIVVELSNSG HGVTMKVPRS
121 EARPSLIGGM LPNAYMFDNI HCHWGKDDIM GSEHFINGES YSMECHMVTY NSKYKDLNDA
181 LSNKDDQDSV VVYTWFYKVK NEDNMKLQQF LNYIPHIRDN GTKINPADQD ALLWFLEPIP
241 ANKEYFAYPG SLTTSPFTPS VTFIILPVPV GISRNQLAVL RTMMDNHNPS KPVLTNRREI
301 QTLNKRPIVC VKGTIQMTAR LIQAEATNDV
```

Active site   Zinc binding site   Signal peptide   **Region: Alpha-carbonic anhydrase (91-312aa)**

**B**

| Position | Amino acid | Replaced amino acid |
|---|---|---|
| 111 | His | Asp |
| 139 | Asn | His |
| 141 | His | Asp |
| 143 | His | Asp |
| 153 | Glu | His |
| 166 | His | Asp |
| 253 | Thr | Asn |

**Figure S2. Characteristics of LssaCA.**

A.Amino acid sequence of LssaCA. The signal peptide is underlined in red. Catalytically active residues are shown in green, and zinc binding sites in blue. The alpha-carbonic anhydrase domain spans residues 91-312.

B.Summary of amino acid substations in the LssaCA mutant. Seven catalytically active residues were replaced with the indicated amino acids to generate an enzymatically inactive mutant.

### Figure S3

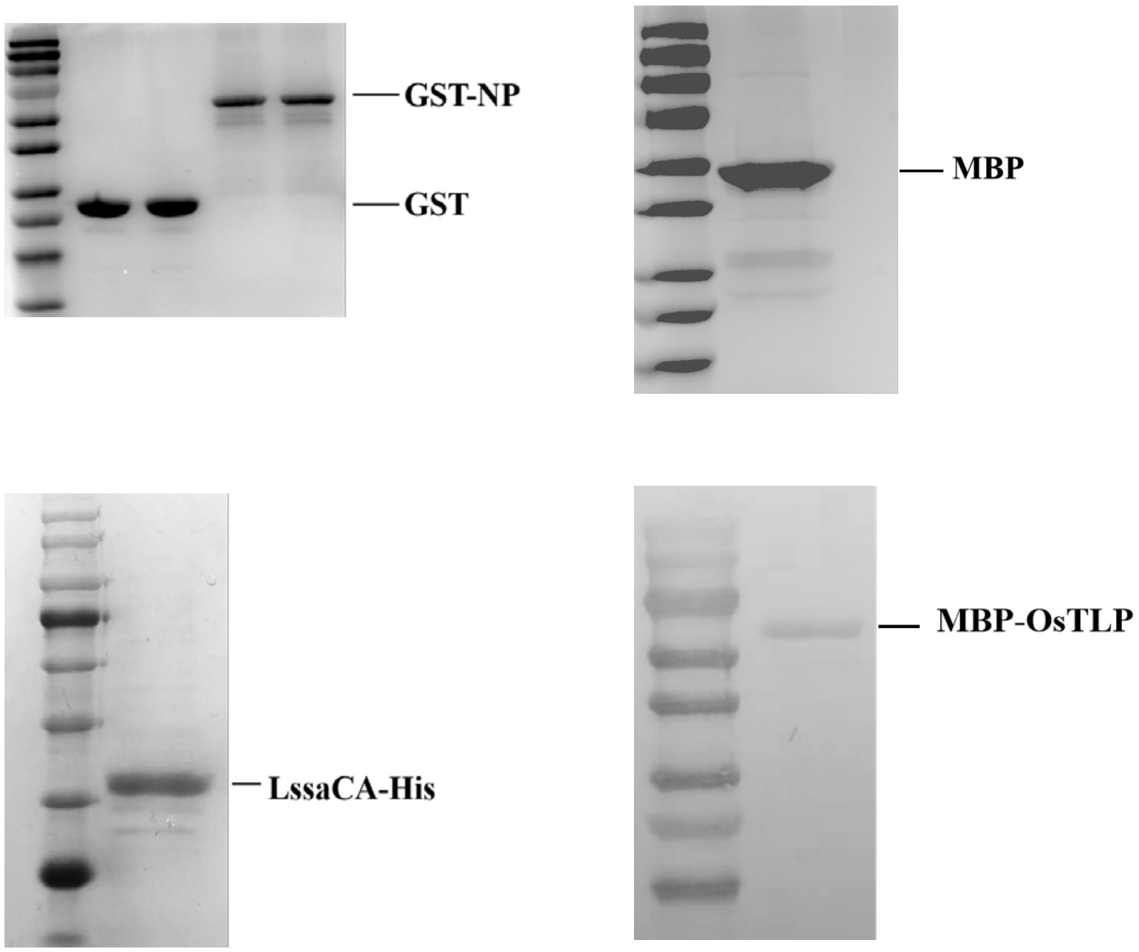


**Figure S3. SDS-PAGE analysis of purified recombinant proteins used in Microscale thermophoresis (MST) assays.**

Protein bands were visualized by Coomassie Brilliant Blue staining.

#### Figure S4

**A**

```
1    ATGTCACAGTGCCCAACTCATGTCGCTCTCCGGTTGCTTGCTCTGCTCTTCCTGCTTCCAGCTGCGTGGTCCGCGACGTTC
1     M  S  Q  C  P  T  H  V  A  L  R  L  L  A  L  L  F  L  L  P  A  A  W  S  A  T  F

82   ACGATGACCAACAATTGCGGCTACACGGTGTGGCCTGGGCTGCTGTCGGGTGCCGGCACGGCGCCGCTGTCGACGACAGGG
28    T  M  T  N  N  C  G  Y  T  V  W  P  G  L  L  S  G  A  G  T  A  P  L  S  T  T  G

163  TTCGCGCTGGCGCACGGCGCGTCGGCGACGGTGGACGCGCCGGCGAGCTGGTCGGGGCGCATGTGGGCGCGCACGCTCTGC
55    F  A  L  A  H  G  A  S  A  T  V  D  A  P  A  S  W  S  G  R  M  W  A  R  T  L  C

244  GCCGAGGACGCGACAGGCAAGTTCACCTGCGCCACGGGGGACTGTGGCTCGGGTGGCATCCAGTGCAACGGCGGTGGCGCG
82    A  E  D  A  T  G  K  F  T  C  A  T  G  D  C  G  S  G  G  I  Q  C  N  G  G  G  A

325  GCGCCGCCCGCCACGCTCATGGAGTTCACGCTCGACGGCTCCGGCGGCATGGACTTCTTCGACGTGAGCCTCGTCGACGGG
109   A  P  P  A  T  L  M  E  F  T  L  D  G  S  G  G  M  D  F  F  D  V  S  L  V  D  G

406  TATAACCTGCCCATGATCATCGTGCCGCAGGGCGGGGGCGCCGCAGCGCCGGCCGGAAGTGGCGGCGGCAGCGGCGGCAAG
136   Y  N  L  P  M  I  I  V  P  Q  G  G  G  A  A  A  P  A  G  S  G  G  G  S  G  G  K

487  TGCATGGCCACGGGCTGCCTCGTCGACCTAAACGGCGCGTGCCCGGCCGACCTGAGGGTCATGGCGGCGAGCACCGGCACC
163   C  M  A  T  G  C  L  V  D  L  N  G  A  C  P  A  D  L  R  V  M  A  A  S  T  G  T

568  GGCGCCGCCGCTCCCGGCGGGGGGCCCGTGGCTTGCCGCAGCGCGTGCGAGGCGTTCGGGTCGCCGCAGTACTGCTGCAGC
190   G  A  A  A  P  G  G  G  P  V  A  C  R  S  A  C  E  A  F  G  S  P  Q  Y  C  C  S

649  GGCGCGTACGGGAACCCGAACACCTGCCGGCCGTCGACCTACTCCCAGTTCTTCAAGAACGCGTGCCCTCGCGCCTACAGC
217   G  A  Y  G  N  P  N  T  C  R  P  S  T  Y  S  Q  F  F  K  N  A  C  P  R  A  Y  S

730  TACGCGTACGACGACTCCACCTCCACATTCACCTGCACCGCCGGCACCAACTACGCCATCACCTTCTGCCCAAGCACCACC
244   Y  A  Y  D  D  S  T  S  T  F  T  C  T  A  G  T  N  Y  A  I  T  F  C  P  S  T  T

811  AGGTGTGAGATCATTCAGACGAGCCAGGAGAACTATTCAACCAACGGCCACCCAGCAGCGCGGCAACGGCTAGGCGGGAGG
271   R  C  E  I  I  Q  T  S  Q  E  N  Y  S  T  N  G  H  P  A  A  R  Q  R  L  G  G  R

892  AATCTGTGCCGCGGCGTGGGTGCGTCTCCCCCCTCCAAGCACAGGACGACAAAAGCTGAGGATGACACCAAGCCTGCACTG
298   N  L  C  R  G  V  G  A  S  P  P  S  K  H  R  T  T  K  A  E  D  D  T  K  P  A  L

973  GCTTTCTAG
325   A  F  *
```

**Figure S4. Nucleotide and deduced amino acid sequence of OsTLP**

The predicted signal peptide is underlined. Conserved thaumatin-like protein sequences are shown in bold.

#### Figure S5

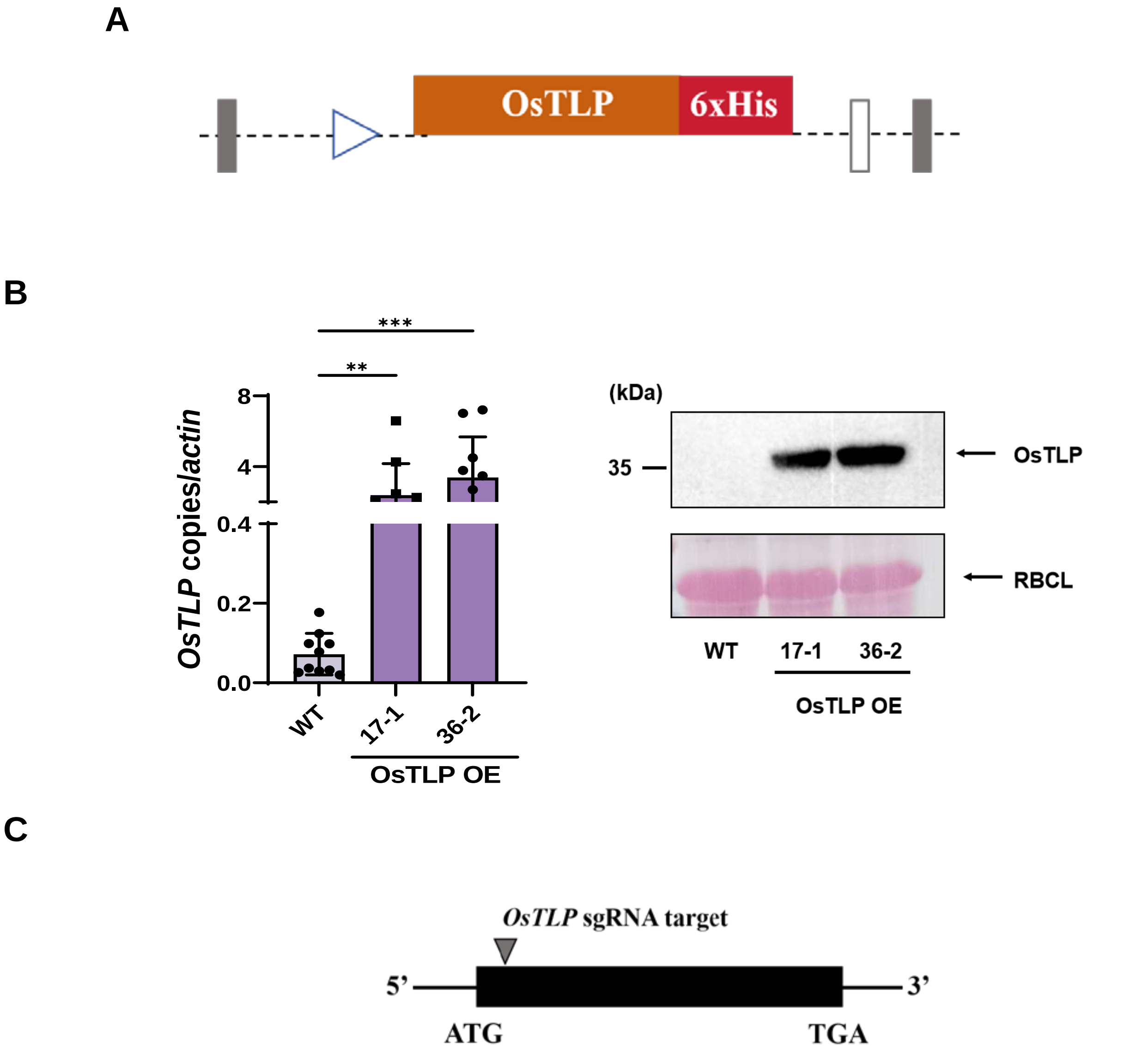


| Line | OsTLP Sequence | Editing |
|---|---|---|
| WT | G G A C T G T G G C T C G G G T G G C A T C | / |
| *ostlp#3* | G G A C T G T G G C T - G G G T G G C A T C | -1 |
| *ostlp#5* | G G A C T G T G G C - - G G G T G G C A T C | -2 |

**Figure S5. Generation of OsTLP-overexpressing and knockout rice lines.**

A.Schematic representation of OsTLP overexpression vector construction using the pCAMBIA1300 backbone, driven by CaMV 35S promoter.
B.RT-qPCR and western blot analyses of OsTLP expression in transgenic lines (OsTLP OE) compared to wild type (WT). OsTLP protein was detected using an anti-His antibody.
C.Summary of mutations in ostlp knockout lines generated via CRISPR/Cas9. Exons are shown as black boxes, with start (ATG) and stop (TGA) codons labeled. The sgRNA target site is marked with a triangle.

#### Figure S6

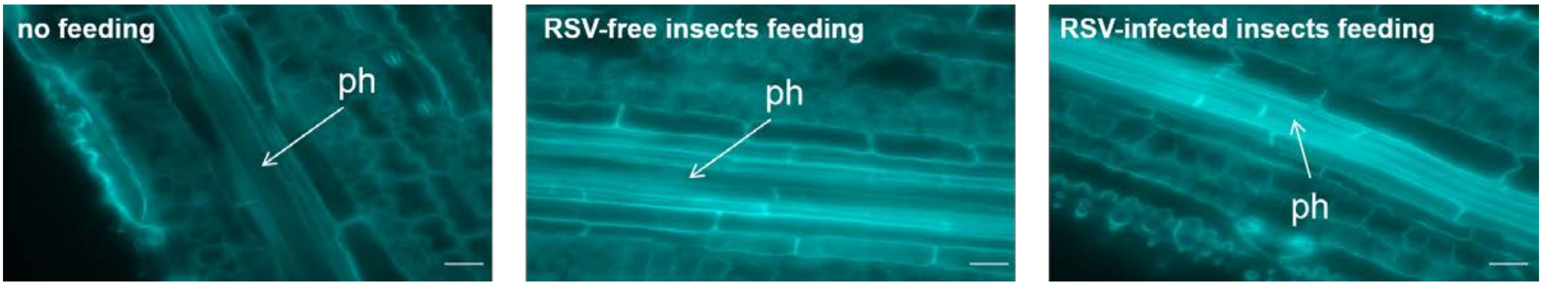


**Figure S6. Detection of callose deposition following *L. striatellus* feeding.**

Bright blue fluorescence indicates callose deposition at feeding sites in longitudinal leaf cross-sections. Samples were collected from plants that were unfed, fed on by RSV-free insects, or fed on by RSV-infected *L. striatellus*. Observation were made using fluorescence microscopy. Abbreviations: xy, xylem; ph, phloem. Scale bars: 10 μm.

### Figure S7

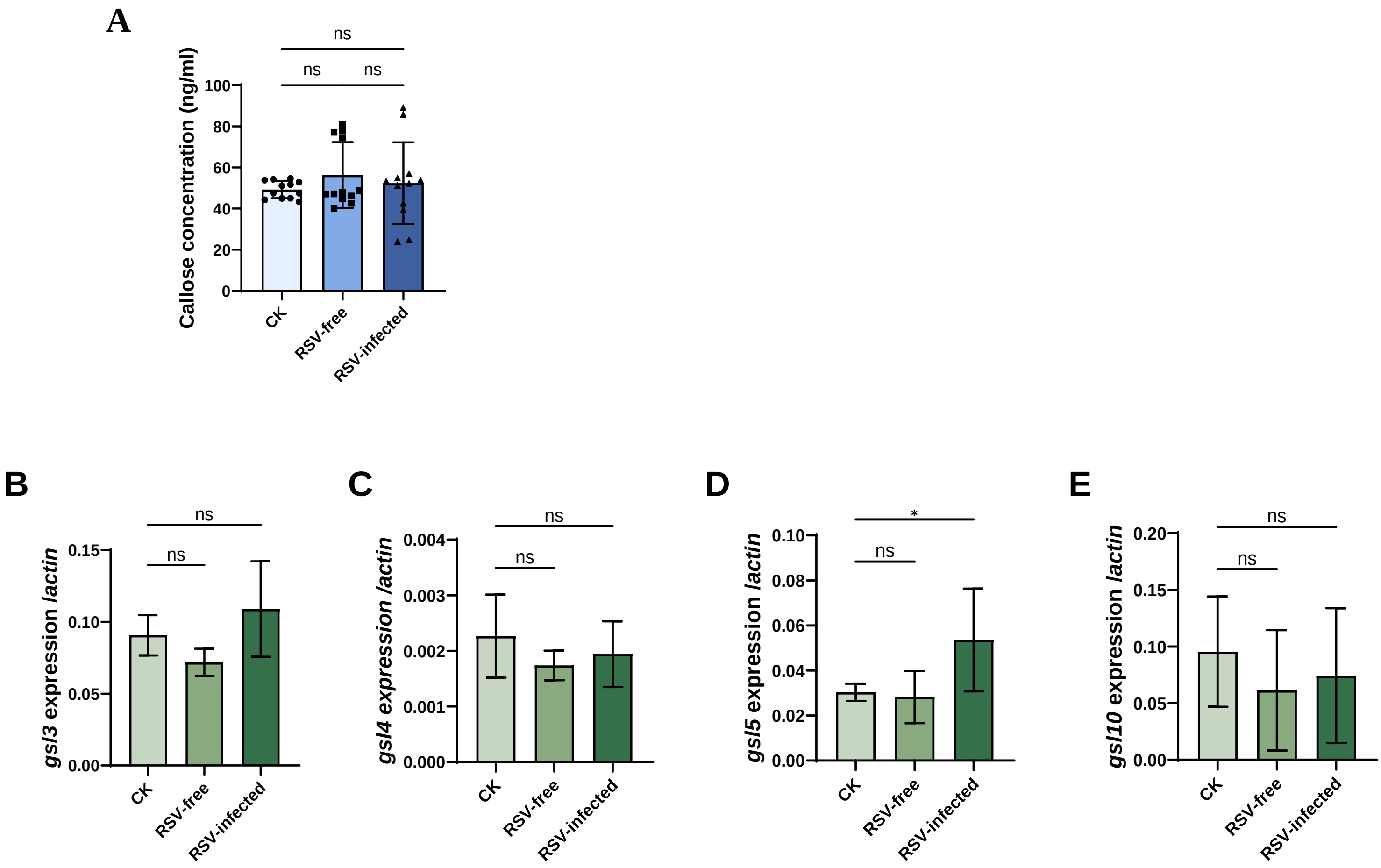


**Figure S7. Callose deposition analysis in non-feeding areas**

A.Callose concentration quantified by ELISA in leaf areas 1 cm away from insect feeding sites. CK: control (non-fed plants).
B-E. RT-qPCR analysis of callose synthase gene expression in non-fed areas. Insects were allowed to feed for 24 h prior to RNA extraction. ns, not significant; *, $p < 0.05$.

**Figure S8**

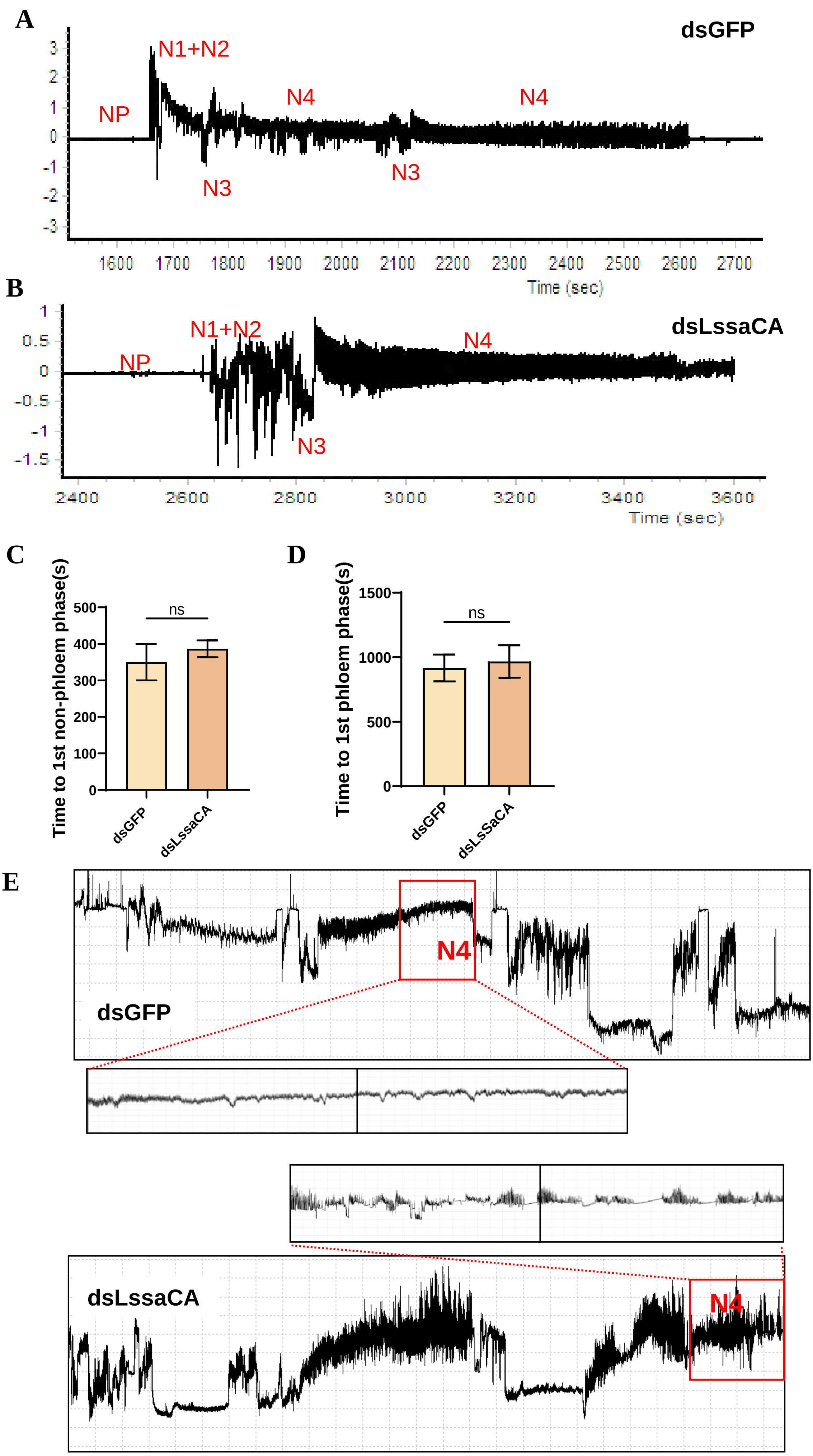
A
dsGFP
N1+N2
NP
N4
N4
N3
N3
Time (sec)
B
dsLssaCA
N1+N2
NP
N4
N3
Time (sec)
C
Time to 1st non-phloem phase(s)
ns
dsGFP
dsLssaCA
D
Time to 1st phloem phase(s)
ns
dsGFP
dsLsSaCA
E
dsGFP
N4
dsLssaCA
N4

**Figure S8. Electrical penetration graph (EPG) to show LssaCA effects on *L. striatellus* feeding behaviors.**

A-B. EPG recordings of *L. striatellus* feeding on rice plants. Waveforms were categorized into non-penetration (NP), penetration (N1, N2, N3), and phloem feeding (N4a & N4b) phases. A, dsGFP-treated insects; B, dsLssaCA-treated insects.
C-D. Quantitative analysis of the time to first non-phloem phase (C) and time to first phloem feeding (D). Mean and SD were calculated from six independent EPG recordings. ns, not significant.
E. EPG recordings showing the continuity of sap ingestion. Red rectangles indicate a 200-second waveform recordings. The N4 waveform of dsLssaCA-treated SBPHs was occasionally interrupted for brief periods.

### Table S1. Yeast two-hybrid screening for rice proteins interacting with LssaCA.

| Yeast Two-Hybrid Screening for Rice Proteins Interacting with LssaCA | |
|---|---|
| **Group ID** | **Protein description** |
| 1 | superoxide dismutase-like protein |
| 2 | thaumatin-like protein 1b isoform X1 |
| 3 | mediator of RNA polymerase II transcription subunit 30 |
| 4 | Bowman-Birk type bran trypsin inhibitor |

**Table S2. List of primers used in this study.**

| Primer name | Primer sequence (5'-3') |
|---|---|
| LssaCA-T7F | TAATACGACTCACTATAGGTACCAAGAAGTGAGGCGAGAC |
| LssaCA-T7R | TAATACGACTCACTATAGGAGAACCTGGGTATGCGAAGTA |
| GFP-T7F | TAATACGACTCACTATAGGATGGTAGATCTGACTAGTAA |
| GFP-T7R | TAATACGACTCACTATAGGCTAGTCATCTGCACCTTCTG |
| LssaCA-q-F | CAGCAGTCGCCTATCGACATT |
| LssaCA-q-R | TCGCCTCACTTCTTGGTACTTTC |
| actin-q-F | AGGAAGGCTGGAAGAGGACC |
| actin-q-R | CGGGAAATTGTGAGGGACAT |
| EF2-q-F | GTCTCCACGGATGGGCTTT |
| EF2-q-R | ATCTTGAATTTCTCGGCATACATTT |
| NP-q-F | GATGCGTTGTCTTACCTGACTGC |
| NP-q-R | CACTATCCCATACCTCGACACCA |
| OsGSL3-q-F | TGGCAAGCGACCACATAG |
| OsGSL3-q-R | AGACCTTAGCACGGACTG |
| OsGSL4-q-F | TGCTTTGAGTGGCACAAGAC |
| OsGSL4-q-R | CTGAAACTCGGAGACGAAGG |
| OsGSL5-q-F | GTGGTGTCCCTGCTATGA |
| OsGSL5-q-R | GTTGTTTGCTATTCTCCC |
| OsGSL10-q-F | TGGCAGCTGTGTTCTTTACG |
| OsGSL10-q-R | CCCATCCGGAAGGTAAGAAT |
| OsTLP-F | GAGAACACGGGGGACTCTAGAATGTCACAGTGCCCAACTCATG |
| OsTLP-R | ACCATGGTGGCTAGCCCGGATCCGAAAGCCAGTGCAGGCTTGG |
| LssaCA-F-BamHI | CATGGAGGCCGAATTCATGGAGCTGTTATTCCACATA |
| LssaCA-R-EcoRI | GCAGGTCGACGGATCCTACATCATTGGTTGCCTCGGCTTG |
| OsTLP-F-1300 | GAGAACACGGGGGACTCTAGAATGTCACAGTGCCCAACTCATG |
| OsTLP-R-1300 | ACCATGGTGGCTAGCCCGGATCCGAAAGCCAGTGCAGGCTTGG |
| OsTLP-MBP-F | GAGGGAAGGATTTCAGAATTCATGTCACAGTGCCCAACTCATG |
| OsTLP-MBP-R | CAGTGCCAAGCTTGCCTGCAGGAAAGCCAGTGCAGGCTTGG |
| OsTLP-pDBleu-F | ATCGTCGAGGTCGACCCCCGGGATGTCACAGTGCCCAACTCATG |
| OsTLP-pDBleu-R | CGGCCGCACTAGTTGCCATGGGATCATGGCGGCAACGAG |
| LssaCA-pPC86-F | AGAGGGTGGGTCGACCCCCGGGATGGAGCTGTTATTCCACATAAGCA |
| LssaCA-pPC86-R | CGCACTAGTAGATCTGGAATTCTACATCATTGGTTGCCTCGGC |
